# Dimethylindanoylindole isomers revealed different pharmacological profiles at CB1 and CB2 cannabinoid receptors

**DOI:** 10.1101/2025.09.16.676463

**Authors:** Soo Jung Oh, Ruiru Guo, Chrisopher Lucaj, Kh Tanvir Ahmed, Gregory B. Dudley, Hideaki Yano

## Abstract

Synthetic cannabinoid receptor agonists (SCRAs) represent one of the most rapidly growing classes of novel psychoactive substances (NPS) and exhibit higher efficacy and potency at CB1 and CB2 cannabinoid receptors (CB1 and CB2) compared to Δ^9^-tetrahydrocannabinol, contributing to distinct adverse effects absent in cannabis use. The indole-based SCRAs GBD-002 and GBD-003, incorporating 2,2-dimethylindane, differ by a single substituent position yet display markedly different receptor binding profiles. We evaluated both the GBD-002/GBD-003 pair and an analogous structural isomer pair using bioluminescence resonance energy transfer (BRET) assays to assess key CB1 and CB2 transducer pathways. Site-directed mutagenesis characterized contributions of non-conserved residues between CB1 and CB2 and aromatic residues on CB1 transmembrane helix 2 (TM2). This study provided insights into the molecular determinants of cannabinoid receptor selectivity and efficacy.

## Introduction

The endocannabinoid system (ECS) is a crucial neuromodulatory network that regulates diverse neurological and neuropsychiatric processes, including mood, movement, reward, appetite, and pain perception ^1–5^. It has become an important target for therapeutic development in conditions such as obesity and chronic pain ^4,6,7^. The ECS signals primarily through two G protein-coupled receptors (GPCRs): CB1 and CB2, among others ^8,9^. Both receptors couple to inhibitory Gα_i/o_ proteins and undergo kinase-mediated phosphorylation, which recruits β-arrestin and leads to receptor desensitization and internalization ^10^.

CB1 is one of the most abundant GPCRs in the brain and plays a central role in modulating neural activity, with its effects varying by neuronal subtype and brain region ^11–13^. In contrast, although CB2 expression is also reported in the brain, it is predominantly expressed in the peripheral immune system, where it acts as a key regulator of immune and inflammatory responses ^14,15^. CB1 and CB2 form the core signaling components of the ECS.

SCRAs, often conceptualized as comprising head, core, and tail moieties, were originally developed as research tools to probe ECS signaling and assess pharmacological potential. However, many early SCRAs were diverted for illicit use, likely due to their synthetic accessibility and initial lack of regulation^16,17^. These compounds bind to CB1 and CB2 and are associated with adverse effects, particularly through CB1 activation ^18^. For example, the naphthoylindole JWH-018 and various analogues emerged in smokable synthetic cannabinoid (SSC) products such as “K2” and “Spice”, prompting regulatory controls that render these compounds impractical for continued use in legitimate pharmacological research.

We recently developed a series of novel 1,3-disubstituted indole-based cannabinoids featuring a more structurally complex and synthetically challenging 2,2-dimethylindanoylindole scaffold, for unrestricted academic research of the ECS ^19,20^. Among these, regioisomers GBD-002 and GBD-003 display markedly different receptor binding profiles. GBD-002 exhibits little selectivity between CB1 and CB2, while GBD-003 is CB2-selective, highlighting molecular interactions that may underlie receptor subtype discrimination ^19^.

CB1 and CB2 are seven-transmembrane (TM) receptors that share a high degree of sequence homology, especially within the TM regions that form the orthosteric binding site ^21^. This similarity may be why many cannabinoids, including Δ^9^-tetrahydrocannabinol (Δ^9^-THC), can bind both receptors with similar affinity^18,22^. While recent advances in cryo-electron microscopy have yielded high-resolution structures of both receptors in active and inactive states, these static snapshots offer limited insight into the dynamic mechanisms that underlie receptor selectivity and function ^23–26^. Understanding the molecular determinants of receptor selectivity, particularly in cases such as GBD-002 and GBD-003, where regioisomers exhibit markedly different pharmacological profiles, can provide insight into how subtle chemical features are differentially recognized by CB1 and CB2.

We hypothesized that non-conserved residues within the orthosteric binding pocket may influence both functional potency and efficacy by differential interactions with ligand headgroup positioning. We investigated the structure–activity relationships (SAR) of GBD-002 and GBD-003 at CB1 and CB2 and used site-directed mutagenesis to characterize key binding site residues that contribute to the CB2 selectivity exhibited by GBD-003 relative to its structural isomer, GBD-002.

## Results

### Isomers with analogous position in head group display similar trends in functional efficacy across CB1 and CB2

We previously demonstrated that the two dimethyl-indane regioisomers, GBD-002 and GBD-003, exhibit distinct binding profiles. In binding assays, GBD-002 showed little to no selectivity between CB1 and CB2 receptors, whereas GBD-003 displayed CB2 selectivity, with a selectivity index of 7.9 ^19^. We then investigated how the orientation of GBD-003’s head group influences functional properties of G protein and β-arrestin recruitment.

To that end, we examined both the GBD-002/GBD-003 pair and an additional structural isomer pair with a different head group but analogous positional variation: JWH-018 and JWH-018 2′-naphthyl isomer (JWH-018 isomer). This allowed us to assess whether receptor selectivity is driven by the specific substituent or by head group attachment position. The efficacy and potency of all four ligands were evaluated at CB1 and CB2 receptors using bioluminescence resonance energy transfer assay (BRET) based biosensors to monitor (1) receptor interaction with Gα_i1_ subunit (Gα_i1_ engagement), (2) receptor recruitment of β-arrestin 2 (β-arrestin 2 recruitment), (3) Gα_i1_ interaction with Gβγ (Gα_i1_ activation), and (4) Gα_oA_ interaction with Gβγ (Gα_oA_ activation). These modes of assays would provide a direct measurement of Gα_i1_ and β-arrestin 2 coupling and would not be influenced by off target receptor activation.

In the CB1 Gα_i1_ engagement assay, GBD-002 demonstrated approximately 60% higher efficacy and an increase in potency compared to its isomer GBD-003 (Figure 1A, Table 1). JWH-018 showed greater efficacy and potency than its isomer, but with a smaller difference compared to the GBD isomer pair. These isomeric trends were consistent in the CB1 β-arrestin 2 recruitment assay (Figure 1C). Similar trends were observed in the CB1 Gα_i1_ activation and Gα_oA_ activation assays (Supplementary Figure S1).

**Table 1.** Efficacy and potency of CB1 and CB2 Gα_i1_ engagement and β-arrestin 2 recruitment assays a.

|  | CB1 |  | CB2 |  |
| --- | --- | --- | --- | --- |
| | $G\alpha_{i1}$<br>Engagement | β-arrestin 2<br>Recruitment | $G\alpha_{i1}$<br>Engagement | β-arrestin 2<br>Recruitment |
| | $E_{\max}$ (%) | | | |
| CP55940 | 99.4 ± 3.0 | 99.8 ± 2.1 | 99.8 ± 2.7 | 99.7 ± 2.7 |
| GBD-002 <sup>b</sup> | 116.4 ± 4.9**** | 163.3 ± 6.6**** | 78.7 ± 8.5 | 91.8 ± 7.3 |
| GBD-003 | 55.1 ± 5.4 | 53.7 ± 7.7 | 75.3 ± 5.2 | 80.8 ± 5.4 |
| JWH-018 <sup>b</sup> | 139.6 ± 5.6** | 167.8 ± 4.7**** | 73.5 ± 3.9 | 75.1 ± 4.4 |
| JWH-018<br>isomer | 110.7 ± 5.0 | 98.4 ± 5.3 | 84.5 ± 6.5 | 91.8 ± 4.6 |
| | $pEC_{50}$ (M) | | | |
| CP55940 | 8.03 ± 0.09 | 7.78 ± 0.06 | 8.76 ± 0.10 | 8.13 ± 0.09 |
| GBD-002 <sup>b</sup> | 7.16 ± 0.10**** | 6.20 ± 0.08*** | 5.86 ± 0.20 | 5.92 ± 0.14 |
| GBD-003 | 5.89 ± 0.16 | 5.44 ± 0.19 | 6.35 ± 0.21 | 6.29 ± 0.14 |
| JWH-018 <sup>b</sup> | 7.61 ± 0.12* | 6.96 ± 0.07* | 8.42 ± 0.19** | 7.68 ± 0.16 |
| JWH-018<br>isomer | 6.99 ± 0.11 | 6.43 ± 0.12 | 7.35 ± 0.23 | 7.40 ± 0.15 |

<sup>a</sup> Mean ± SEM values for $E_{\max}$ and $pEC_{50}$ are reported. $E_{\max}$ is normalized against CP55940.
<sup>b</sup> GBD-002 and JWH-018 have \* $p < 0.05$ , \*\*\*\* $p < 0.0001$ denoting differences relative to their respective isomers, GBD-003 and JWH-018 naphthyl, by one-way ANOVA followed by Šidák's post hoc test.

**Table 2.** Reciprocal substitutions of non-conserved residues in CB1 and CB2 binding pocket. c.

| CB1 mutant receptors: Gα<sub>i1</sub> Engagement |  |  |  |  |  |
| --- | --- | --- | --- | --- | --- |
|  | WT | L193I | I267L | L359V | Triple Mutations |
|  | E<sub>max</sub> (%) |  |  |  |  |
| CP55940 | 99.4 ± 3.0 | 98.2 ± 2.2 | 77.6 ± 2.5*** | 122.3 ± 3.3*** | 160.5 ± 1.6**** |
| GBD-002 | 116.4 ± 4.9 | 83.6 ± 3.0** | 67.5 ± 4.1**** | 124.4 ± 4.1 | 140.5 ± 2.4 |
| GBD-003 | 55.1 ± 5.4 | 56.2 ± 4.3 | 26.9 ± 2.9* | 40.3 ± 4.7 | 88.9 ± 1.9** |
| JWH-018 | 139.6 ± 5.6 | 91.8 ± 2.1**** | 75.4 ± 4.5**** | 126.7 ± 3.4 | 148.7 ± 1.5 |
| JWH-018 isomer | 110.7 ± 5.0 | 106.1 ± 2.8 | 72.6 ± 4.0**** | 102.4 ± 4.5 | 148.6 ± 2.6**** |
|  | pEC<sub>50</sub> (M) |  |  |  |  |
| CP55940 | 8.03 ± 0.09 | 7.31 ± 0.06 | 7.92 ± 0.10 | 7.75 ± 0.08 | 7.76 ± 0.03 |
| GBD-002 | 7.16 ± 0.10 | 7.36 ± 0.10 | 6.94 ± 0.15 | 7.19 ± 0.09 | 7.56 ± 0.05 |
| GBD-003 | 5.89 ± 0.16 | 6.15 ± 0.14 | 6.78 ± 0.26 | 6.27 ± 0.23 | 6.55 ± 0.05 |
| JWH-018 | 7.61 ± 0.12 | 7.88 ± 0.07 | 7.38 ± 0.17 | 7.71 ± 0.08 | 7.89 ± 0.03 |
| JWH-018 isomer | 6.99 ± 0.11 | 6.95 ± 0.06 | 6.87 ± 0.13 | 7.07 ± 0.11 | 7.09 ± 0.04 |

| CB2 mutant receptors: Gα<sub>i1</sub> Engagement |  |  |  |  |  |
| --- | --- | --- | --- | --- | --- |
|  | WT | I110L | L182I | V261L | Triple Mutations |
|  | E<sub>max</sub> (%) |  |  |  |  |
| CP55940 | 99.8 ± 2.7 | 83.1 ± 2.8*** | 32.2 ± 2.5**** | 33.8 ± 2.6**** | 3.0 ± 2.0**** |
| GBD-002 | 78.7 ± 8.5 | 66.8 ± 6.9 | 17.6 ± 2.7**** | 24.7 ± 3.0**** | -1.9 ± 2.0**** |
| GBD-003 | 75.3 ± 5.2 | 74.2 ± 7.0 | 26.3 ± 3.6*** | 32.4 ± 12.5** | -5.4 ± 1.7**** |
| JWH-018 | 73.5 ± 3.9 | 58.3 ± 5.9 | 15.1 ± 3.6**** | 29.9 ± 4.1**** | -0.4 ± 2.3**** |
| JWH-018 isomer | 84.5 ± 6.5 | 66.5 ± 6.5 | 23.5 ± 3.9**** | 64.5 ± 9.7 | 10.8 ± 2.2**** |
|  | pEC<sub>50</sub> (M) |  |  |  |  |
| CP55940 | 8.76 ± 0.10 | 8.81 ± 0.13 | 8.30 ± 0.27 | 9.32 ± 0.35 | 8.23 ± 9.48 |
| GBD-002 | 5.86 ± 0.20 | 5.34 ± 0.40 | 9.35 ± 0.53 | 8.10 ± 0.46 | 8.73 ± 6.11 |
| GBD-003 | 6.35 ± 0.21 | 5.16 ± 0.46 | 7.35 ± 0.42 | 5.55 ± 0.60 | 11.05 ± 2.40* |
| JWH-018 | 8.42 ± 0.19 | 7.96 ± 0.35 | 8.38 ± 0.84 | 9.23 ± 0.59 | 8.74 ± 3.29 |
| JWH-018 isomer | 7.35 ± 0.23 | 7.31 ± 0.27 | 7.37 ± 0.49 | 6.08 ± 0.30 | 8.74 ± 0.98 |

<sup>c</sup> \**p* < 0.05, \*\*\*\**p* < 0.0001 denote differences in WT to mutant compound efficacies, determined by one-way ANOVA with Dunnett's post hoc test.

**Figure 1.**
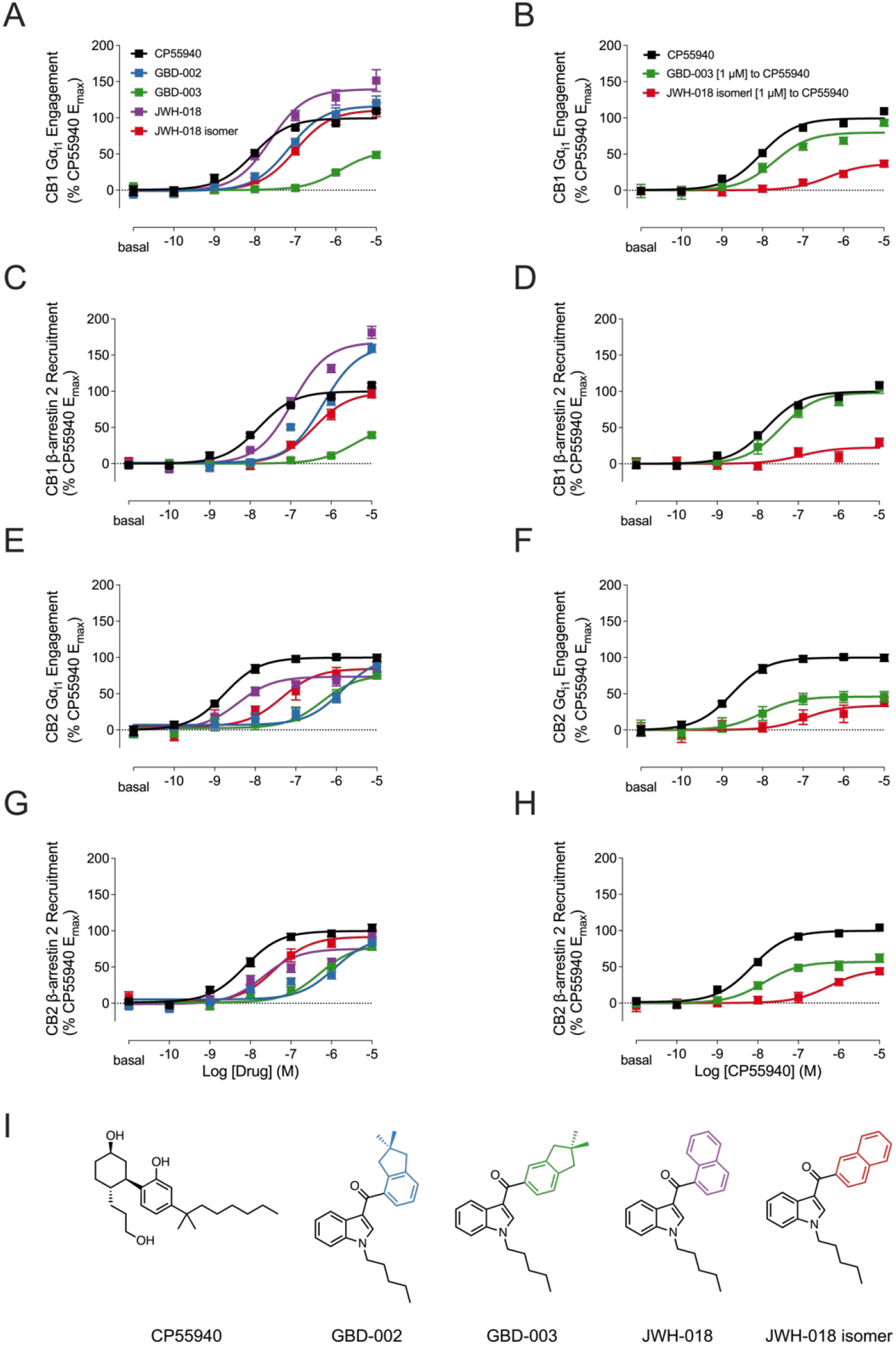
BRET concentration-response curves of CB1 Gα_i1_ engagement (A), CB1 β-arrestin 2 recruitment (C), CB2 Gα_i1_ engagement (E), and CB2 β-arrestin 2 recruitment (G) measured in response to CP55940, GBD-002, GBD-003, JWH-018, and JWH-018 isomer. Vehicle, 1 µM GBD-003, and JWH-018 isomer were pre-incubated before the addition of CP55940 (B, D, F, and H). Data presented as means ± SEM of n=4 experiments. (I) Chemical structures shown from left to right: CP55940, GBD-002, GBD-003, JWH-018, and JWH-018 isomer.

In contrast, CB2 Gα_i1_ engagement assay revealed no significant differences between GBD-002 and GBD-003, though GBD-003 trended toward three-fold higher potency (Figure 1E, Table 1). Compared to CB1, GBD-002 exhibited reduced efficacy and potency at CB2, whereas GBD-003 showed increased activity, supporting its CB2 selectivity. These findings were mirrored in the CB2 β-arrestin 2 recruitment assay, where both GBD compounds displayed similar efficacies and potencies with no significant differences (Figure 1G).

The JWH isomers showed a general trend of reduced efficacy but increased potency in both CB2 Gα_i1_ engagement and β-arrestin 2 recruitment assays compared to the CB1 assays (Figure 1E, G). Notably, JWH-018 had significantly higher potency than JWH-018 isomer in the CB2 Gα_i1_ engagement assay but did not differ in potency in the β-arrestin 2 assay.

To further investigate the increased efficacy of GBD-003 in the CB2 assay compared to CB1, we performed a pre-stimulation assay in which cells were first stimulated with 1µM GBD-002 or GBD-003 followed by addition of the full agonist CP55940 (Figure 1B, D, F, H). This approach allowed us to assess the functional efficacy of partial agonists prior to being superseded by the high affinity and efficacy of CP55940. In this competitive context, the partial agonists (GBD-003 and JWH-018 isomer) act as antagonists, occupying the binding site and producing submaximal responses.

In both CB1 Gα_i1_ engagement and β-arrestin 2 assays, pre-stimulation of GBD-003 elicited minimal response and only reduced CP55940’s maximal efficacy by ∼10%, consistent with its low efficacy at CB1. In contrast, JWH-018 isomer significantly reduced CP55940 efficacy by 70% in both assays, reflecting higher efficacy. At CB2, however, GBD-003 reduced CP55940 efficacy by ∼50% in both Gα_i1_ engagement and β-arrestin 2 assays, while JWH-018 isomer maintained a ∼60% reduction. These results corroborate results testing GBD-003 and JWH-018 isomer efficacy.

Overall, our results demonstrate that the head group positioning influences receptor efficacy. GBD-003 displayed higher efficacy at CB2 compared to CB1, whereas GBD-002 showed the opposite trend. For the JWH isomers, both showed lower efficacy at CB2, but JWH-018 isomer exhibited higher efficacy than JWH-018, reversing their relative efficacies at CB1.

### Reciprocal substitutions of non-conserved residues between CB1 and CB2 influence efficacy but not selectivity

We next investigated how differences in binding between the isomeric pairs contribute to their potency and efficacy profiles using site-directed mutagenesis in CB1 and CB2. Molecular dynamics simulations revealed two key non-conserved residues in CB1 and CB2 that influence the binding of GBD-002 and GBD-003 ^19^. In CB1, residues L193^3.29^ and L359^6.51^ stabilize GBD-002 and form a hydrogen bond with S383^7.39 19^. However, in the case of GBD-003, the positioning of the indane ring introduces steric strain, shifting the ligand toward extracellular loop 2(ECL2) and disrupting the hydrogen bond with S383^7.39 19^. In contrast, CB2 contains shorter residues at the corresponding positions, I110^3.32^ and V261^6.51^, which allow both ligands to rotate within the binding site while maintaining hydrogen bonding with S285^7.39 19^. Furthermore, ECL2 is a divergent region between CB1 and CB2. It has been proposed that CB1 residue I267^ECL2^, located within ECL2, contributes to ligand selectivity by creating additional space within the CB1 binding pocket, whereas the corresponding bulkier residue in CB2 may cause steric hindrance ^24,27^.

We then tested whether reciprocal substitutions of these non-conserved residues could recreate the opposite receptor’s ligand-binding characteristics. For CB1, we made the following mutations: L193I^3.29^, I267L^ECL2^, L359V^6.51^, and a triple mutant containing all three substitutions (CB1 triple mutant). For CB2, we generated I110L^3.32^, L182I^ECL2^, V261L^6.51^, and a corresponding triple mutant (CB2 triple mutant). Triple mutants would assess how the combined effects of the three residues influence ligand efficacy and potency. BRET ratios were normalized to the WT receptor response to CP55940 to enable comparison of each isomer’s efficacy relative to the wild-type baseline. This allowed us to assess how individual mutations altered the efficacy of each isomer and to compare those changes across mutations.

The CB1 L193I^3.29^ mutation resulted in a significant reduction in efficacy for the CB1-selective agonists GBD-002 (∼30%) and JWH-018 (∼50%) compared to the wild-type (WT) receptor (Figure 2C). In contrast, GBD-003 and JWH-018 isomer showed minimal reductions in efficacy. The CB1 I267L^ECL2^mutation caused a general decrease in efficacy for all ligands, and this effect was particularly pronounced for GBD-002 (∼50%) and JWH-018 (∼65%), with nearly twice the decrease observed for their respective isomers GBD-003 (∼30%) and JWH-018 isomer (∼40%) (Figure 2D).

**Figure 2.**
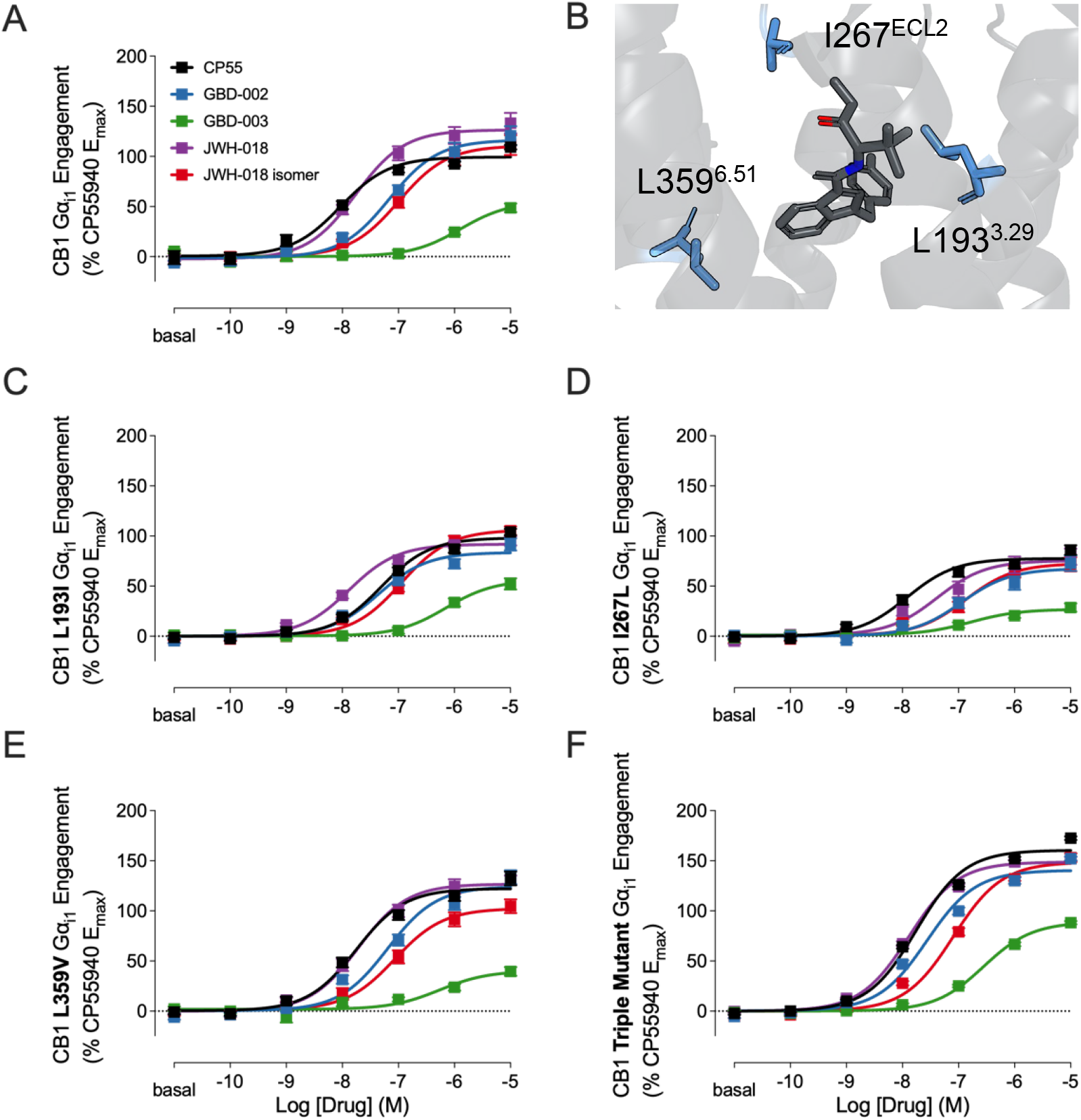
Gα_i1_ engagement BRET concentration-response curves of CB1 (A) WT compared to (C) L193I, (D) I267L, (E) L359V, and (F) triple mutant for CP55940, GBD-002, GBD-003, JWH-018, and JWH-018 isomer. Data presented as means ± SEM of n=4 experiments. (B) CB1 non-conserved residues (blue) interacting with MDMB-FUBINACA (dark grey) PDB: 6N4B.

Interestingly, the CB1 L359V^6.51^ mutation showed a trend toward increased efficacy for CP55940 and GBD-002, and a decrease for GBD-003, JWH-018, and the JWH-018 isomer (Figure 2E). In the CB1 triple mutant (L193I^3.29^, I267L^ECL2^, and L359V^6.51^), all compounds displayed increased efficacies (Figure 2F). Notably, GBD-003 (∼35%) and JWH-018 isomer (∼40%) showed statistically higher efficacy compared to the WT receptor. Potency changes across all CB1 mutants were not statistically significant. Similar changes in efficacy and potency were observed in the CB1 triple mutant β-arrestin 2 assay (Supplementary Figure S2A).

The reductions in efficacy for GBD-002 and JWH-018 at L193I^3.29^ and I267L^ECL2^, alongside the increased efficacy of GBD-003 and the JWH-018 isomer in the triple mutant, suggest that they partially mimic the CB2 receptor’s binding pocket in efficacy.

For the CB2 receptor mutants, the I110L^3.32^ substitution resulted in a decrease in efficacy across all ligands (∼1% to 20%) (Figure 3C). The CB2 L182I^ECL2^ mutation caused a significant reduction in efficacy for all compounds (∼50 to 70%), with GBD-003 exhibiting a slightly smaller decrease (Figure 3D). Similarly, the CB2 V261L^6.51^ mutation significantly reduced efficacy for all compounds except the JWH-018 isomer, which showed only a ∼20% decrease (Figure 3E). Interestingly, the CB2 triple mutant (I110L^3.32^, L182I^ECL2^, and V261L^6.51^) abolished agonist efficacy for all tested compounds (Figure 3F). No significant changes in potency were observed for any CB2 mutant relative to WT. In line with the Gα_i1_ result, no drug-induced BRET was observed in the CB2 triple mutant β-arrestin 2 assay (Supplementary Figure S2B).

**Figure 3.**
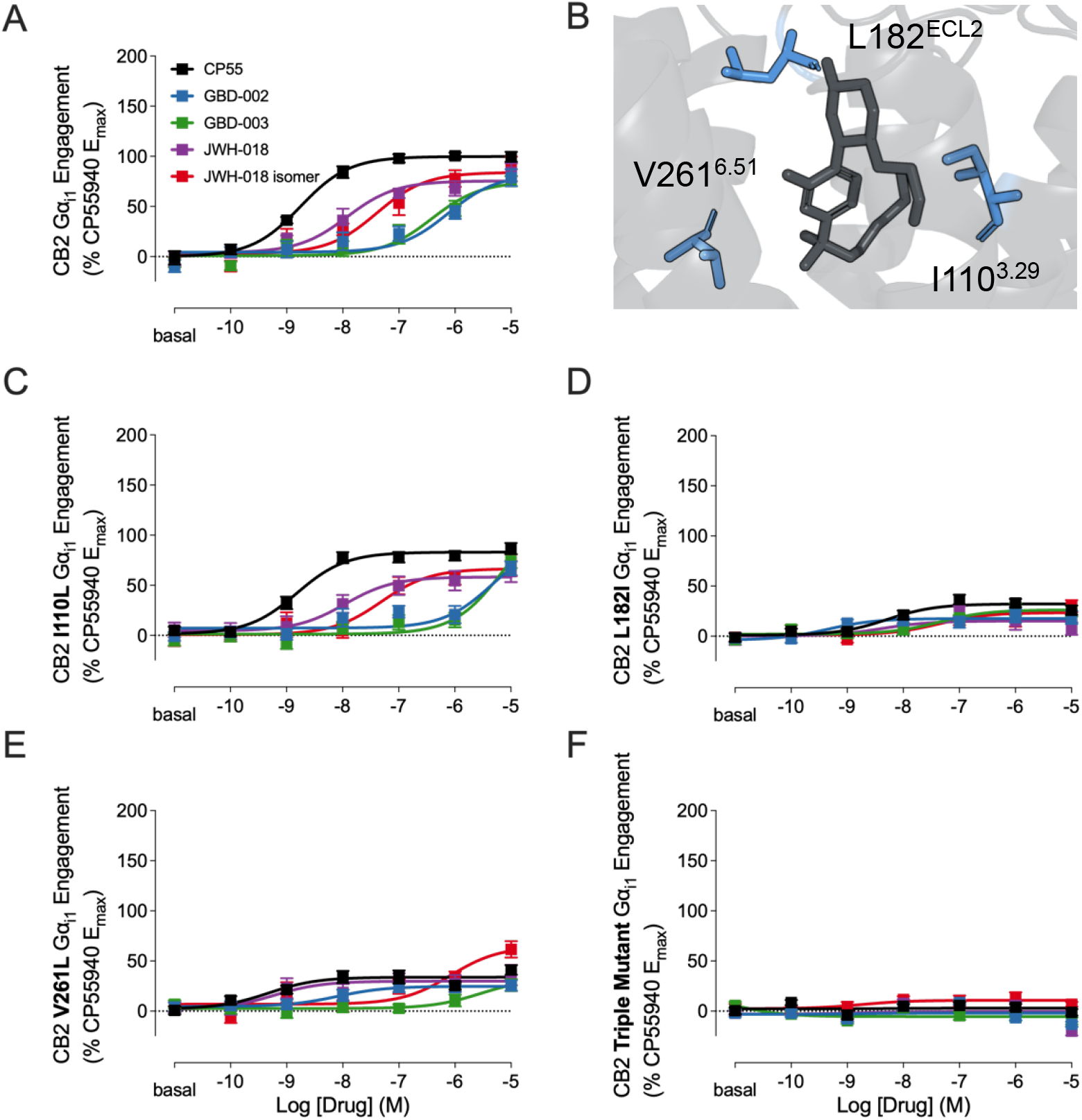
Gα_i1_ engagement BRET concentration-response curves of CB2 (A) WT compared to (C) I110L, (D) L182I, (E) V261L, and (F) triple mutant for CP55940, GBD-002, GBD-003, JWH-018, and JWH-018 isomer. Data presented as means ± SEM of n=4 experiments. (B) CB2 non-conserved residues (blue) interacting with CP55940 (dark grey) PDB: 8GUR.

Overall, CB2 mutants primarily exhibited substantial reductions in efficacy, in contrast to the reciprocal mutations in CB1 with partially CB2-like efficacy. A possible explanation may be the structural difference between the two receptors: the CB1 binding pocket is generally larger and more flexible, whereas the CB2 pocket is narrower and more rigid ^28^. As a result, amino acid substitutions in CB2 may cause effects beyond ligand interactions, disrupting receptor dynamics due to the compact binding site ^24,28^.

### Alanine substitution of TM2 residues reveals their significant role in modulating ligand efficacy

In comparison to CB2, CB1 receptor activation induces a structural rearrangement in transmembrane helix 2 (TM2), which has been proposed to play a key role in modulating ligand efficacy ^24,29,30^. This shift repositions aromatic residues F170^2.57^, F174^2.61^, F177^2.64^, and H178^2.65^ toward the headgroup of MDMB-FUBINACA, a synthetic cannabinoid receptor agonist structurally similar to the GBD and JWH compounds ^24^. Given that efficacy was the primary differentiating factor between the isomeric ligand pairs, we examined whether variations in headgroup positioning influenced interactions with these TM2 residues. To test this, we performed alanine-scanning mutagenesis and evaluated changes in ligand efficacy.

The F170A^2.57^ mutation caused a significant reduction in E_max_ for all compounds (Figure 4C). The JWH-018 and JWH-018 isomer were most affected, exhibiting ∼100% and ∼80% decreases, respectively, while GBD-002 and GBD-003 showed more moderate reductions (∼60% and ∼50%). Interestingly, the F174A^2.61^ mutation led to a 32% increase in efficacy for GBD-002, while GBD-003 showed a downward trend (Figure 4D). In contrast, both JWH compounds exhibited significant decreases in efficacy, indicating that F174^2.61^ may play a more selective role in modulating efficacy for JWH ligands and GBD-003, but not GBD-002. The F177A^2.64^ mutation resulted in a total reduction in efficacy across all compounds (Figure 4E). Finally, the H178A^2.65^ mutation caused a substantial decrease in efficacy for the JWH compounds (∼80%), while having a more moderate effect on the GBD compounds (∼35%) (Figure 4F).

**Figure 4.**
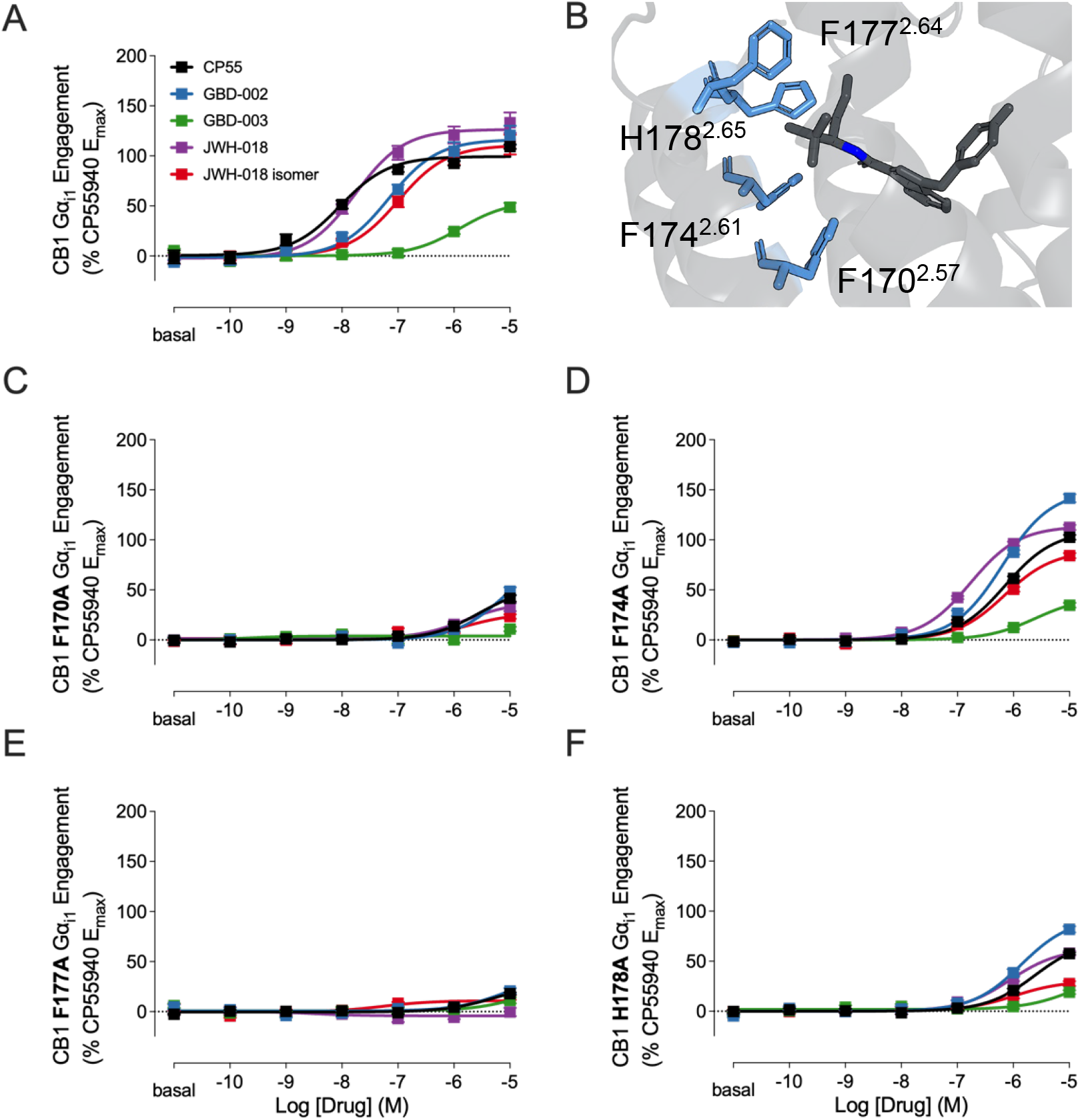
Gα_i1_ engagement BRET concentration-response curves of CB1 (A) WT compared to TM2 mutants (C) F170A, (D) F174A, (E) F177A, and (F) H178A for CP55940, GBD-002, GBD-003, JWH-018, and JWH-018 isomer. Data presented as means ± SEM of n=4 experiments. CB1 TM2 residues (blue) interacting with MDMB-FUBINACA (dark grey) PDB: 6N4B.

For all TM2 mutations, alanine substitution of each residue reduced overall ligand efficacy while preserving the rank order of ligand responses, suggesting that TM2 modulates efficacy independently of headgroup chemical composition and or spatial orientation. Together, these findings indicate that the TM2 helix plays a significant role in modulating ligand efficacy.

## Discussion

To investigate how the position of dimethyl-indane isomers GBD-002 and GBD-003 influences CB2 selectivity, we utilized BRET to monitor the direct coupling of wild-type and mutant CB1 and CB2 receptors with Gα_i/o_ and β-arrestin 2. Our findings demonstrate that the orientation of dimethyl-indane influences potency at CB1 but not CB2, while modulating efficacy at both receptors. Single amino acid substitutions of non-conserved orthosteric residues between CB1 and CB2 revealed that introducing

CB2 residues into CB1 partially recapitulated CB2-like efficacy but not potency, whereas the reverse CB1-to-CB2 substitutions had no effect. Additionally, mutagenesis of CB1 TM2 residues demonstrated their significant role in modulating ligand efficacy.

Contrary to our initial hypothesis that substituting non-conserved orthosteric residues between CB1 and CB2 would reproduce both efficacy and potency, only the CB1 triple mutant was able to restore CB2-like efficacy. A recent study using the CB2-selective agonist YL025 proposed that ligand recognition in CB1 and CB2 is driven predominantly by entropic factors, such as binding pocket flexibility, rather than enthalpic contributions from direct ligand–receptor interactions ^28^. In this model, the CB2 binding site is relatively rigid and undergoes minimal conformational change upon activation, favoring interactions with rigid ligands. In contrast, CB1 undergoes more extensive rearrangement upon activation, particularly involving inward movement of TM1 and TM2, making it more dynamic and better suited to accommodate flexible ligands ^24,28^. This entropic explanation may account for why the reciprocal mutations did not result in significant potency shifts due to their conservative changes (e.g., leucine to isoleucine or valine).

Further, triple substitutions of non-conserved orthosteric residues led to striking divergence in receptor activity: introducing CB2 residues into CB1 (CB1 triple mutant) significantly enhanced efficacy, whereas introducing CB1 residues into CB2 (CB2 triple mutant) abolished it. This is consistent with computational studies showing that the antagonist-bound conformation of CB2 resembles the agonist-bound conformation of CB1, with similar pocket volumes and nearly identical residue arrangements ^26^. Our data suggest that while CB1 and CB2 share overall structural homology in their binding pockets, a small number of non-conserved residues function as critical molecular switches for receptor activation. The fact that subtle side-chain substitutions (e.g., leucine to isoleucine or valine) yielded such dramatic effects underscores the sensitivity of the residues of the orthosteric binding pocket to cannabinoid receptor activation and may explain how shifts in ligand head group orientation can differentially influence receptor activation.

This study provided insights into the molecular determinants of cannabinoid receptor selectivity and efficacy. The dimethyl-indane position in GBD compounds influenced both receptor selectivity and efficacy: GBD-002 showed high efficacy at both receptors without selectivity, while GBD-003 exhibited CB2 selectivity with enhanced efficacy. These distinct profiles demonstrate the unique utility of dimethylindane as a versatile headgroup for fine-tuning CB1 and CB2 receptor interaction. Mutagenesis revealed contributions of non-conserved orthosteric residues to CB1 and CB2 activation, while TM2 mutations revealed their importance in modulating CB1 efficacy. Although point mutations may introduce broader conformational effects beyond ligand binding, these results offer a valuable framework for structure-guided drug design leveraging dimethylindane’s modularity and contribute to a mechanistic understanding of how SCRA structural features govern cannabinoid receptor function.

## Methods

### Cell Culture

Human embryonic kidney 293T (HEK-293T) cells were cultured in Dulbecco’s modified Eagle’s medium (DMEM, Corning) supplemented with 10% fetal bovine serum, 2 mM L-glutamine (Corning), and 1% penicillin-streptomycin (Corning). Cells were maintained in 10-cm plates at 37°C with 5% CO_2_.

### Bioluminescence Resonance Energy Transfer (BRET) Assays

HEK-293T cells were transiently transfected with cDNA (10-15 µg per assay) using linear polyethyleneimine hydrochloride (PEI Max ®, Polysciences) at a 1:2 ratio (30 µg). The plasmid weight and transfection ratio were optimized and maintained across BRET assays. The transfection reagent-plasmid mixture was added dropwise to the medium and incubated overnight.

Four BRET Assays were used in this experiment: Gα_i1_ engagement, β-arrestin 2 recruitment, Gα_i1_ activation, and Gα_oA_ activation. For Gα_i1_ engagement, *Renilla* luciferase 8 (RLuc) fused receptor and Venus fused Gα_i1_ served as a resonance energy transfer (RET) pair. Gβ_1_ and Gγ_2_ constructs were co-transfected to avoid depletion of endogenous Gβγ. The β-arrestin 2 recruitment assay utilized an RLuc fused receptor and a β-Arrestin 2 fused Venus as a RET pair. To enhance receptor phosphorylation and facilitate β-Arrestin 2 recruitment, GRK2 constructs were co transfected. Gα_i1_ and Gα_oA_ activation assay used RLuc fused Gα and Venus fused Gγ_2_ as a RET pair. Receptor and Gβ_1_ constructs were co transfected.

### Bioluminescence Resonance Energy Transfer (BRET) Experiment

48 hours post-transfection, cells were washed with PBS, lifted using 5 mM EDTA (Fisher Scientific), collected and resuspended in PBS containing 0.1% glucose, 0.1% BSA, and 200 µM sodium bisulfite. Cells were distributed in white U-bottom 96-well plates, and ligands were added as agonists or antagonists. Agonists were transferred as 10-fold serial dilution in triplicate. Antagonists were pre-incubated with cells at 1 µM for 10 minutes before the addition of the agonist CP55940. Cells were incubated with 5 µM luciferase substrate (coelenterazine H) for 2.5 minutes before agonist transfer. BRET signals were recorded 10 minutes after ligand treatment using the PheraStar FSX plate reader. G protein engagement and dissociation were assessed by BRET ratios (530/480nm) from RLuc and Venus-tagged proteins. Each condition was performed in four to five biological replicates.

### Compounds and Plasmid Constructs

GBD-002 and GBD-003 compounds were prepared as described previously^19^. All other CB1 ligands were obtained from Cayman Chemical (Ann Arbor, Michigan) and dissolved at 10 mM in DMSO. DNA constructs were subcloned in pcDNA3 vectors. Site directed mutagenesis was performed on plasmid constructs encoding *CNR1* and *CNR2* using the Quickchange Site Directed Mutagenesis Kit (Agilent) with the following primers. Mutations were confirmed by GENEWIZ, Azenta Life Sciences.

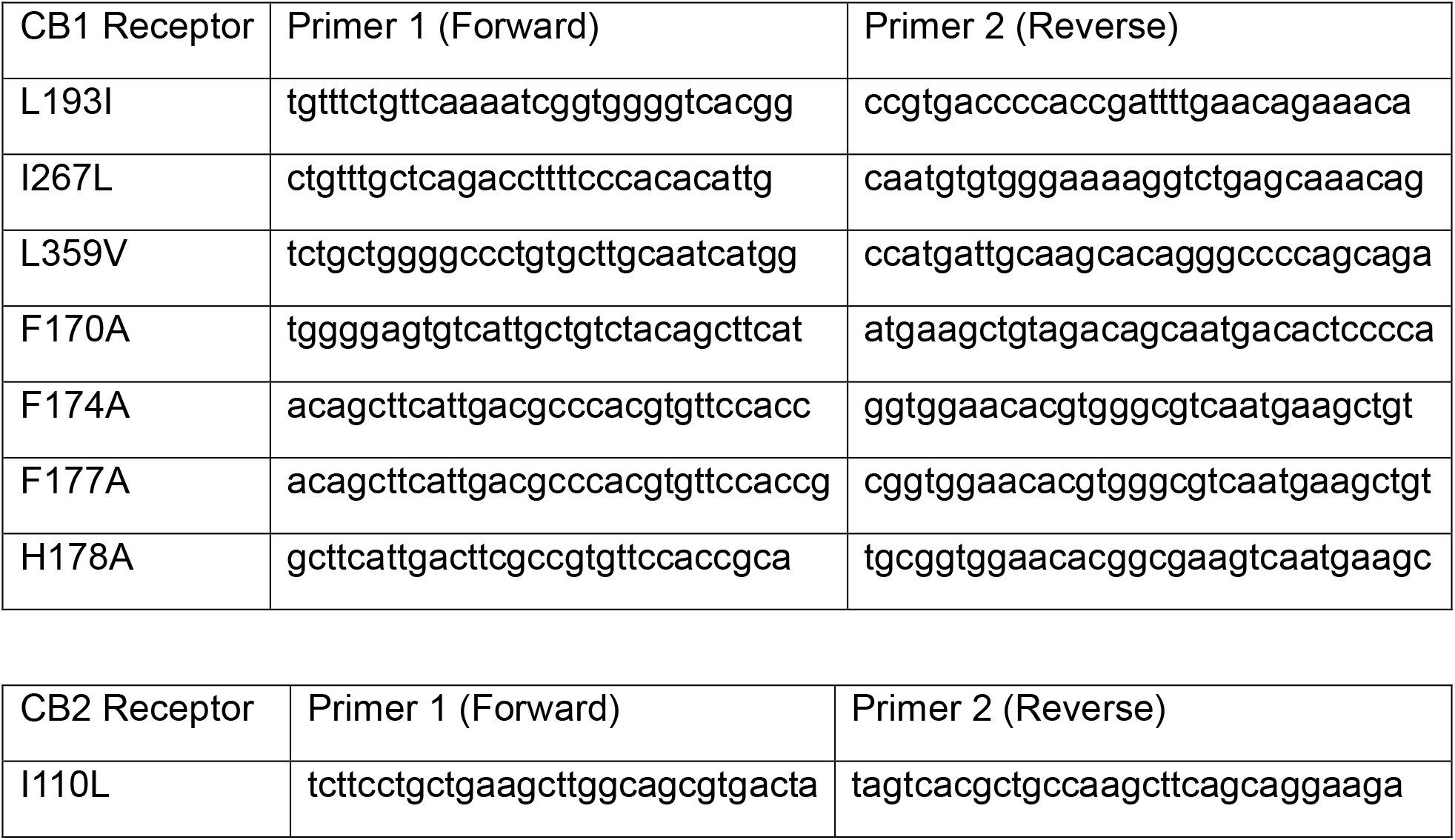
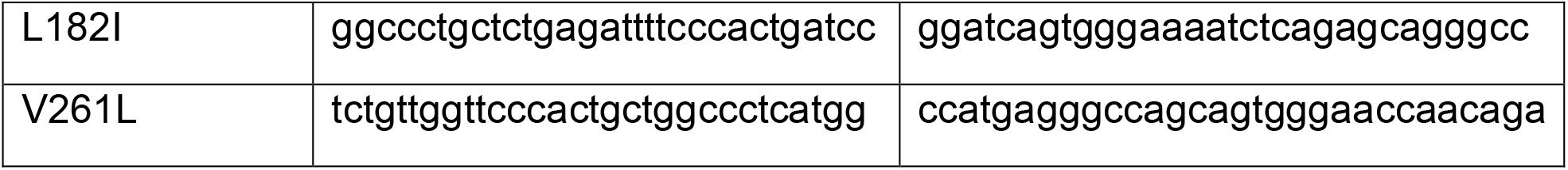

### Statistical Analysis

BRET ratios (530/480 nm) were used to generate four-parameter dose-response curves and analyzed for ANOVA one-way statistical comparison in GraphPad Prism v10. All data were normalized to maximal and minimal BRET signal ratio of WT reference agonist CP55940. Non-linear regression was performed on compiled data to generate E_max_ and pEC_50_.

## Supplemental Figures

**Supplemental Figure 1.**
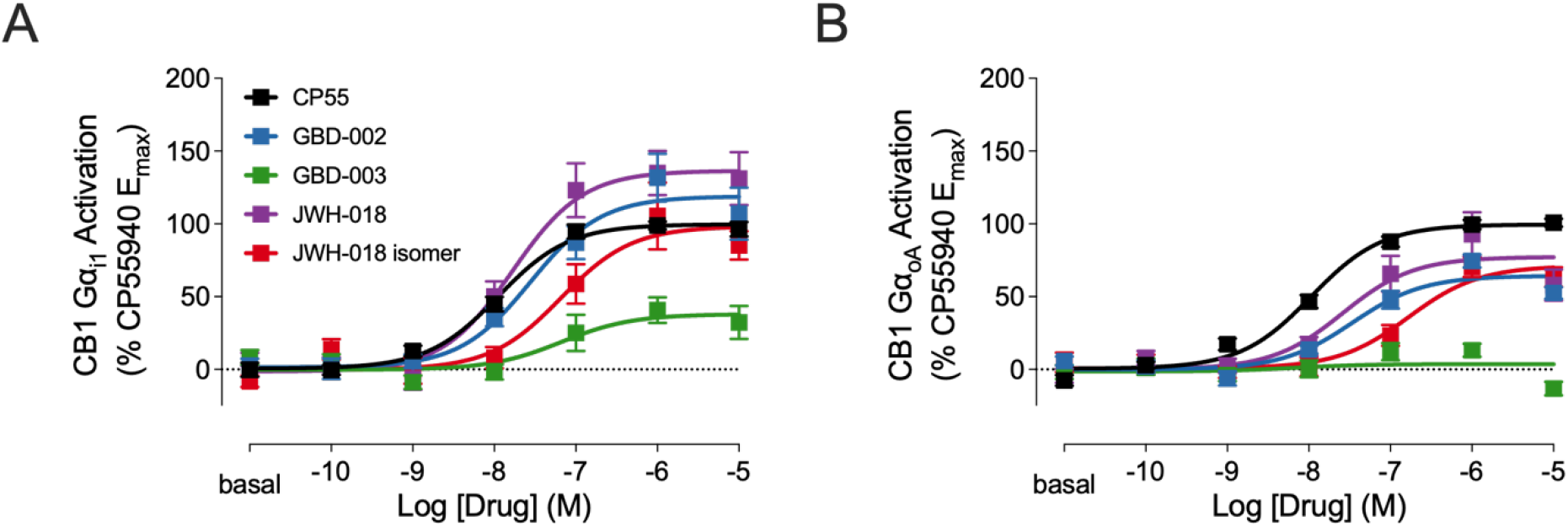
BRET concentration-response curves of (A) CB1 Gα_i1_ interaction with Gβγ (Gα_i1_ activation), and (B) Gα_oA_ interaction with Gβγ (Gα_oA_ activation) measured in response to CP55940, GBD-002, GBD-003, JWH-018, and JWH-018 isomer. Data presented as means ± SEM of n=4 experiments.

**Supplemental Figure 2.**
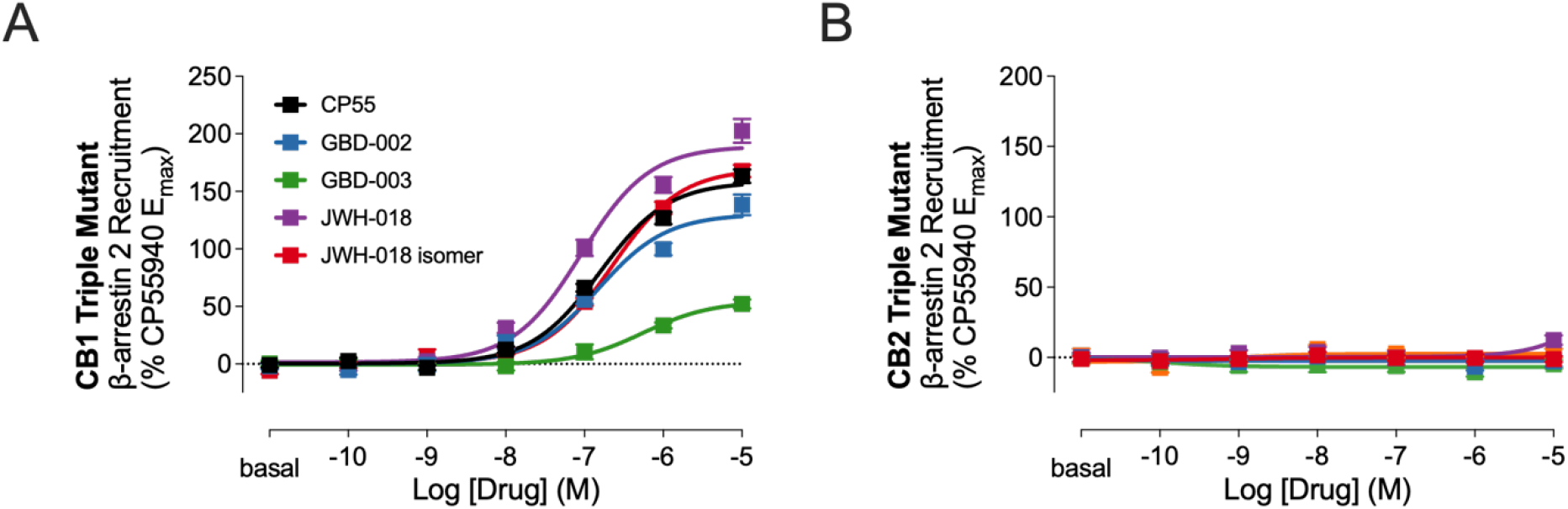
BRET concentration-response curves of β-arrestin 2 recruitment assays for (A) CB1 triple mutant (L193I, I267L, L359V) and (B) CB2 triple mutant (I110L, L182I, V261L) with CP55940, GBD-002, GBD-003, JWH-018, and JWH-018 isomer Data presented as means ± SEM of n=4 experiments.

---

**Supplemental Table 1:**
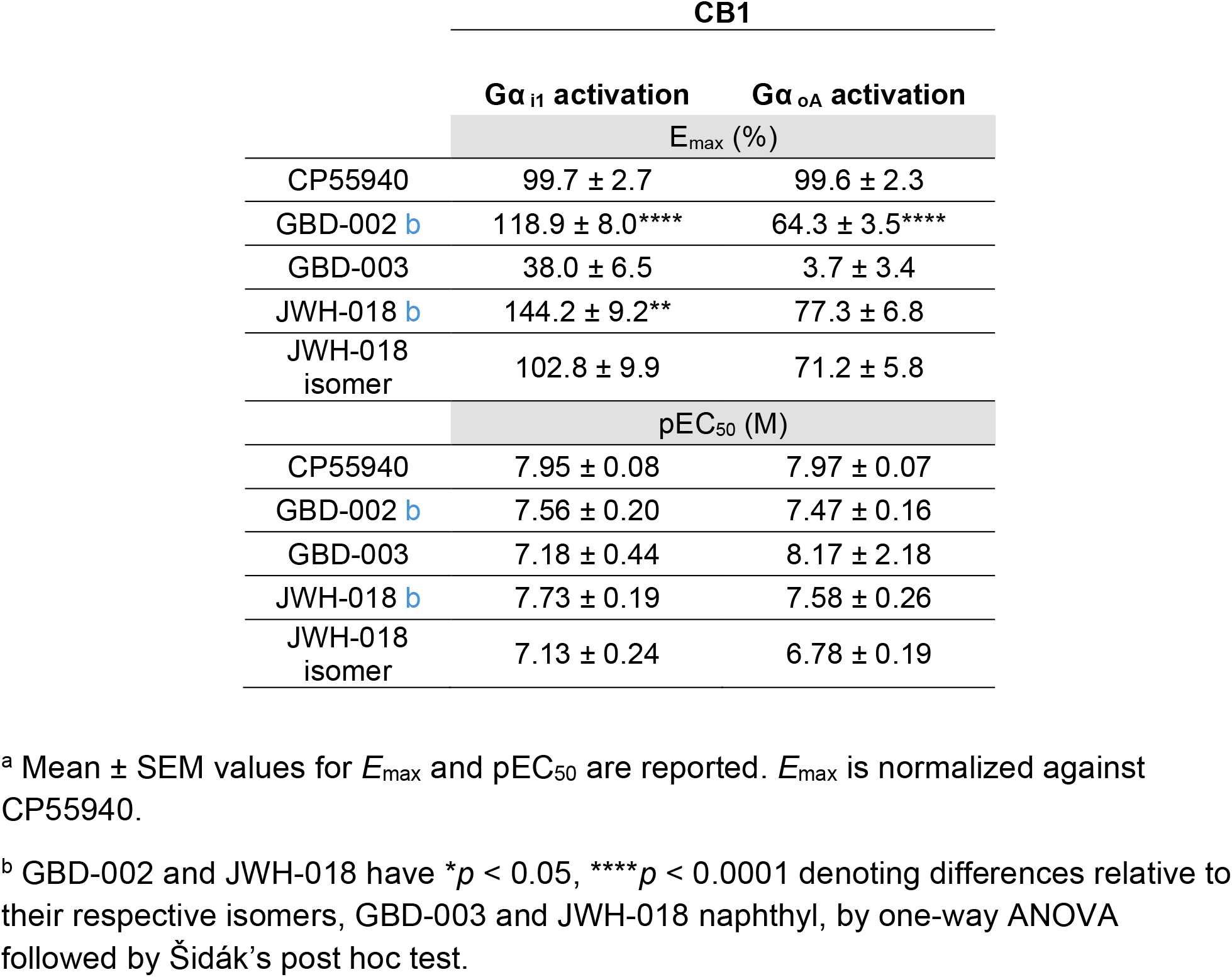
Efficacy and potency of CB1 Gα_i1_ and Gα_oA_ activation a.

| **CB1** |  |  |
| --- | --- | --- |
|  | **$G\alpha_{i1}$ activation** | **$G\alpha_{oA}$ activation** |
|  | **$E_{\max}$ (%)** |  |
| CP55940 | 99.7 ± 2.7 | 99.6 ± 2.3 |
| GBD-002 <sup>b</sup> | 118.9 ± 8.0<sup>****</sup> | 64.3 ± 3.5<sup>****</sup> |
| GBD-003 | 38.0 ± 6.5 | 3.7 ± 3.4 |
| JWH-018 <sup>b</sup> | 144.2 ± 9.2<sup>**</sup> | 77.3 ± 6.8 |
| JWH-018 isomer | 102.8 ± 9.9 | 71.2 ± 5.8 |
|  | **pEC<sub>50</sub> (M)** |  |
| CP55940 | 7.95 ± 0.08 | 7.97 ± 0.07 |
| GBD-002 <sup>b</sup> | 7.56 ± 0.20 | 7.47 ± 0.16 |
| GBD-003 | 7.18 ± 0.44 | 8.17 ± 2.18 |
| JWH-018 <sup>b</sup> | 7.73 ± 0.19 | 7.58 ± 0.26 |
| JWH-018 isomer | 7.13 ± 0.24 | 6.78 ± 0.19 |

<sup>a</sup> Mean ± SEM values for $E_{\max}$ and pEC<sub>50</sub> are reported. $E_{\max}$ is normalized against CP55940.
<sup>b</sup> GBD-002 and JWH-018 have \* $p$ < 0.05, \*\*\*\* $p$ < 0.0001 denoting differences relative to their respective isomers, GBD-003 and JWH-018 naphthyl, by one-way ANOVA followed by Šidák's post hoc test.

**Supplemental Table 2.** Efficacy and potency of CB1 G α_i1_ engagement for TM2 mutants F170, F174, F177, and H178. c.

| CB1 TM2 mutant receptors: G<sub>i1</sub> Engagement |  |  |  |  |  |
| --- | --- | --- | --- | --- | --- |
|  | WT | F170A | F174A | F177A | H178A |
|  | E<sub>max</sub> (%) |  |  |  |  |
| CP55940 | 99.4 ± 3.0 | 53.0 ± 5.6**** | 109.1 ± 2.4 | 18.0 ± 4.7**** | 57.7 ± 5.2**** |
| GBD-002 | 116.4 ± 4.9 | 48.7 ± 3.8**** | 149.1 ± 3.4** | 20.7 ± 11.1**** | 81.9 ± 11.8** |
| GBD-003 | 55.1 ± 5.4 | 4.0 ± 1.4**** | 43.1 ± 4.2 | 11.3 ± 11.7*** | 19.1 ± 7.3** |
| JWH-018 | 139.6 ± 5.6 | 37.7 ± 4.7**** | 113.6 ± 2.0**** | -4.2 ± 1.8**** | 62.6 ± 2.9**** |
| JWH-018 isomer | 110.7 ± 5.0 | 26.9 ± 5.1**** | 89.2 ± 3.2** | 10.9 ± 2.2**** | 30.5 ± 2.6**** |
|  | pEC<sub>50</sub> (M) |  |  |  |  |
| CP55940 | 8.03 ± 0.09 | 5.56 ± 0.15**** | 6.15 ± 0.04**** | 5.30 ± 0.28**** | 5.61 ± 0.08**** |
| GBD-002 | 7.16 ± 0.10 | 5.12 ± 0.25**** | 6.20 ± 0.04 | 5.12 ± 0.64**** | 5.86 ± 0.07** |
| GBD-003 | 5.89 ± 0.16 | 9.77 ± 1.48 | 5.61 ± 0.15 | 4.78 ± 2.76 | 5.02 ± 0.92 |
| JWH-018 | 7.61 ± 0.12 | 5.81 ± 0.21 | 6.77 ± 0.04 | 8.55 ± 1.40 | 6.06 ± 0.08 |
| JWH-018 isomer | 6.99 ± 0.11 | 5.77 ± 0.32** | 6.16 ± 0.07 | 7.26 ± 0.57 | 5.98 ± 0.15* |

<sup>c</sup> \* $p < 0.05$ , \*\*\*\* $p < 0.0001$ denote differences in WT to mutant compound efficacies, determined by one-way ANOVA with Dunnett's post hoc test.

## Notes

### Competing Interest Statement

The authors have declared no competing interest.

